# Prospects for detecting early warning signals in discrete event sequence data: application to epidemiological incidence data

**DOI:** 10.1101/2020.04.02.021576

**Authors:** Emma Southall, Michael J. Tildesley, Louise Dyson

## Abstract

Early warning signals (EWS) identify systems approaching a critical transition, where the system undergoes a sudden change in state. For example, monitoring changes in variance or autocorrelation offers a computationally inexpensive method which can be used in real-time to assess when an infectious disease transitions to elimination. EWS have a promising potential to not only be used to monitor infectious diseases, but also to inform control policies to aid disease elimination. Previously, potential EWS have been identified for prevalence data, however the prevalence of a disease is often not known directly. In this work we identify EWS for incidence data, the standard data type collected by the Centers for Disease Control and Prevention (CDC) or World Health Organization (WHO). We show, through several examples, that EWS calculated on simulated incidence time series data exhibit vastly different behaviours to those previously studied on prevalence data. In particular, the variance displays a decreasing trend on the approach to disease elimination, contrary to that expected from critical slowing down theory; this could lead to unreliable indicators of elimination when calculated on real-world data. We derive analytical predictions which can be generalised for many epidemiological systems, and we support our theory with simulated studies of disease incidence. Additionally, we explore EWS calculated on the rate of incidence over time, a property which can be extracted directly from incidence data. We find that although incidence might not exhibit typical critical slowing down properties before a critical transition, the rate of incidence does, presenting a promising new data type for the application of statistical indicators.

**Author summary:** The threat posed by infectious diseases has a huge impact on our global society. It is therefore critical to monitor infectious diseases as new data become available during control campaigns. One obstacle in observing disease emergence or elimination is understanding what influences noise in the data and how this fluctuates when near to zero cases. The standard data type collected is the number of new cases per day/month/year but mathematical modellers often focus on data such as the total number of infectious people, due to its analytical properties. We have developed a methodology to monitor the standard type of data to inform whether a disease is approaching emergence or disease elimination. We have shown computationally how fluctuations change as disease data get closer towards a tipping point and our insights highlight how these observed changes can be strikingly different when calculated on different types of data.

## Introduction

One of the greatest challenges in society today is the burden of infectious diseases, affecting public health and economic stability all over the world. Infectious diseases disproportionately affect individuals in poverty, with millions of those suffering daily from diseases that are considered eradicable. The potential for eradicating diseases such as polio, guinea worm, measles, mumps or rubella is immense (International Task Force for Disease Elimination, [31]). Even where effective vaccines or treatments exist, disease elimination presents an ongoing challenge. For example, after the establishment of the Global Malaria Eradication Program in 1955 by the World Health Organisation (WHO) it was later abandoned in 1969 due to funding shortages and drug resistance [26], leading to re-emergence of disease in Europe [27]. Assessing when a disease is close enough to elimination to die out without further intervention, thus prompting the end of a control campaign, is a problem of global economic importance. If campaigns are stopped prematurely it can result in disease resurgence and subsequently put control efforts back by decades. Conversely, the threat posed by newly emerging diseases such as SARs, Ebola or the recent corona-virus outbreak COVID-2019 strains available resources, places restrictions on global movement and disrupts the worlds most vulnerable societies. Identifying which newly-emerging diseases will present a global threat, and which will never cause a widespread epidemic is of critical importance.

To overcome the challenges identifying disease elimination or emergence, numerous studies have suggested the use of early warning signals (EWS) [3–8]. EWS are statistics that may be derived from data that change in a predictable way on the approach to a critical threshold. In epidemiology this threshold is commonly described as the point at which the basic reproduction number, *R*_0_, passes through *R*_0_ = 1. A system with *R*_0_ increasing through 1 describes an emerging or endemic disease whereas *R*_0_ decreasing through 1 results in disease elimination. We seek to find EWS to identify when a disease is approaching such a transition. We may identify such statistics using critical slowing down (CSD) theory, which indicates the imminent approach of a threshold, arising from increasing recovery times of perturbations as a system approaches a critical transition [1, 2]. This increase in recovery time occurs because, as the stability of a steady state changes, such as from a disease free state to emergence or from endemic state to elimination, the dominant eigenvalue of the steady state passes through zero. Since the eigenvalue also determines the relaxation time of the system, this recovery time therefore increases as we approach a critical transition.

EWS offer the ability to anticipate a critical transition indirectly in real world noisy time series data, by observing, for example, increasing variance or autocorrelation in the fluctuations around the steady-state [2, 9]. Statistical indicators offer a computationally inexpensive and efficient method for assessing the status of an infectious disease, presenting a simple mechanism for disease surveillance and monitoring of control policies.

The development of EWS is an active area of research in many fields, identifying the statistical signatures of abrupt shifts in many dynamical systems. Studies have applied EWS to historical data or laboratory experiments where a tipping point is known [1, 13, 23]; developed methods for using spatial variation [17, 18], explored the effects of detrending [8, 15] using the ensemble of multiple EWS [12, 13, 30]; and developed understanding of the limitations of EWS [16, 19, 20].

Discrepancies in statistical signatures have been discovered in a variety of historical datasets known to be going through a critical transition: from climate systems to stock markets, to applications with ecological field data [16, 22, 23]. These studies observed unexpected characteristic traits of common EWS, such as identifying a decreasing trend in variance or standard deviation, leading to a discussion on the robustness of indicators. It is therefore highly important to understand analytically how EWS are expected to change on the approach to a critical transition for different data types to avoid any misleading results.

The initial development of EWS in epidemiology focused on prevalence data, producing analytical solutions and numerically testing the capabilities for statistical indicators of emergence and elimination of infectious diseases [5–8]. Analysis of computer simulations of well-studied epidemiological systems have highlighted challenges such as seasonality [6] or detrending of epidemiological time series data [8]. However epidemiological data is typically collected in the form of the number of new infectious cases (incidence data) over a certain period of time (weekly/monthly/yearly). Generally, the exact date of infection or recovery of an individual is not known and therefore the exact number of infectious individuals at each point in time (the prevalence data that has been analysed) is unknown.

Simulation-based studies exploring incidence-type data have suggested that the potential for emergence of an infectious disease can be informed by statistical signatures [3, 4]. These studies represent the first attempts to understand the robustness of some indicators when used with disease emergence incidence data, subject to underreporting and time aggregation. Both studies find that EWS do precede disease emergence even when reporting is low. When the numerical performance of 10 EWS are compared, Brett *et al.* find that the mean and variance perform well unless incidence is subject to a highly overdispersed reporting error and they compare these results with previously studied prevalence results. Theoretical predictions are given for prevalence data, however the analytical behaviour of incidence is not explored.

O’Dea *et al.* [4] incorporate an observation model into a Birth-Death-Immigration (BDI) process to present an analytical study of EWS of disease emergence. This model allows prediction of the behaviour of EWS for dynamics captured by a BDI process but is not suitable for diseases with population-level immunity. O’Dea *et al.* additionally conducted an investigation into reporting errors in incidence-type data by recording the removal of individuals (“death” component in the BDI process). They describe the probability of a case being reported with either a Binomial or Negative Binomial distribution, allowing for over and under-reporting. In contrast to Brett *et al.*, they conclude that the mean, variance and coefficient of variation (CV) are poor indicators since they are sensitive to reporting errors and insensitive to differences between transmission and recovery rates.

In this paper, we advance the current literature to describe generalised signatures of statistical indicators for incidence data, on the approach to a threshold, highlighting the differences between EWS descriptors of incidence and prevalence. Strikingly, our results demonstrate that although EWS of emergence exhibit an increasing variance, a trait associated with CSD and supporting results from Brett *et al.* and O’Dea *et al.*, we demonstrate that variance instead *decreases* on the approach to a disease elimination transition. We find that although incidence data does not undergo the transcritical bifurcation traditionally considered by CSD theory, nevertheless time series trends are still a valuable tool to predict disease elimination. The discrepancy between prevalence and incidence on the one hand, and elimination and emergence on the other, could lead to potential problems in detecting thresholds if the differences are not clearly understood.

We introduce an analytical theory from stochastic processes to address why variance in incidence decreases for disease elimination. We study multiple other indicators of disease elimination predicted by this theory, and compare their responses with stochastic simulations. We also consider the rate of incidence as a measurement that can be extracted from incidence data. Notably, we find that on the approach to a critical transition the rate of incidence exhibits typical CSD signatures which correspond with prevalence data, such as an increasing variance. We present a broad analytical framework for EWS of incidence and rate of incidence for a variety of different disease systems. We explore more intricate systems where elimination is driven by different factors to understand the robustness of this theory. This simple generalised result can be applied to many infectious diseases undergoing emergence or elimination, a promising development for EWS of infectious diseases.

## Methods & Mathematical Theory

In this paper we focus on the application of EWS to disease elimination where there is a limited understanding on how time series statistics of incidence data behaves on the approach towards this threshold. We consider two simple models, where disease elimination is forced with different mechanisms, to explore how EWS of disease elimination behave for prevalence and incidence data. To demonstrate the broadness of our results, we additionally present a comparative case study to the analytical results for emerging diseases by O’Dea *et al.*

In this section we review the following models: SIS model (Susceptible-Infected-Susceptible model, see for example Keeling & Rohani [24]); SIS model with vaccination and SIS model with external force of infection. For each model that we have chosen to investigate, we derive the stochastic differential equations (SDEs) that describe the analytical behaviour of prevalence in these systems. Derivations of the analytical results and calculations of each statistic can be found in the supporting text (S1 Appendix). We present our analysis for incidence data and derive the corresponding statistical indicators. We exploit the well known fact that a counting process can be described by a Poisson process. We apply this result to the field of EWS to incorporate statistical signatures of a Poisson distributed variable to describe the behaviour of the number of new infectious cases in epidemiology.

We verify our analytical results for prevalence and incidence with simulated studies, and compare the contrasting results between prevalence and incidence. We measure the change in trend of multiple statistical indicators using the Kendall’s Tau score which gives an indication of an increasing or decreasing trend.

## Prevalence Theory

### Model 1: SIS with reducing transmission

We begin with a simple example of a system that is approaching elimination from an existing endemic state of *I*. We consider an SIS model where the effective contact rate *β* acts as the control parameter. Effective reduction of *β* can be induced by public health campaigns (such as washing hands or improving food hygiene) and through social distancing (such as school closure). By decreasing *β*(*t*) in time, it slowly forces 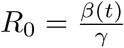 through the critical transition at *R*_0_ = 1. The model transitions are shown in the following schematic with transition probabilities given in Table 1,

**Table 1.**
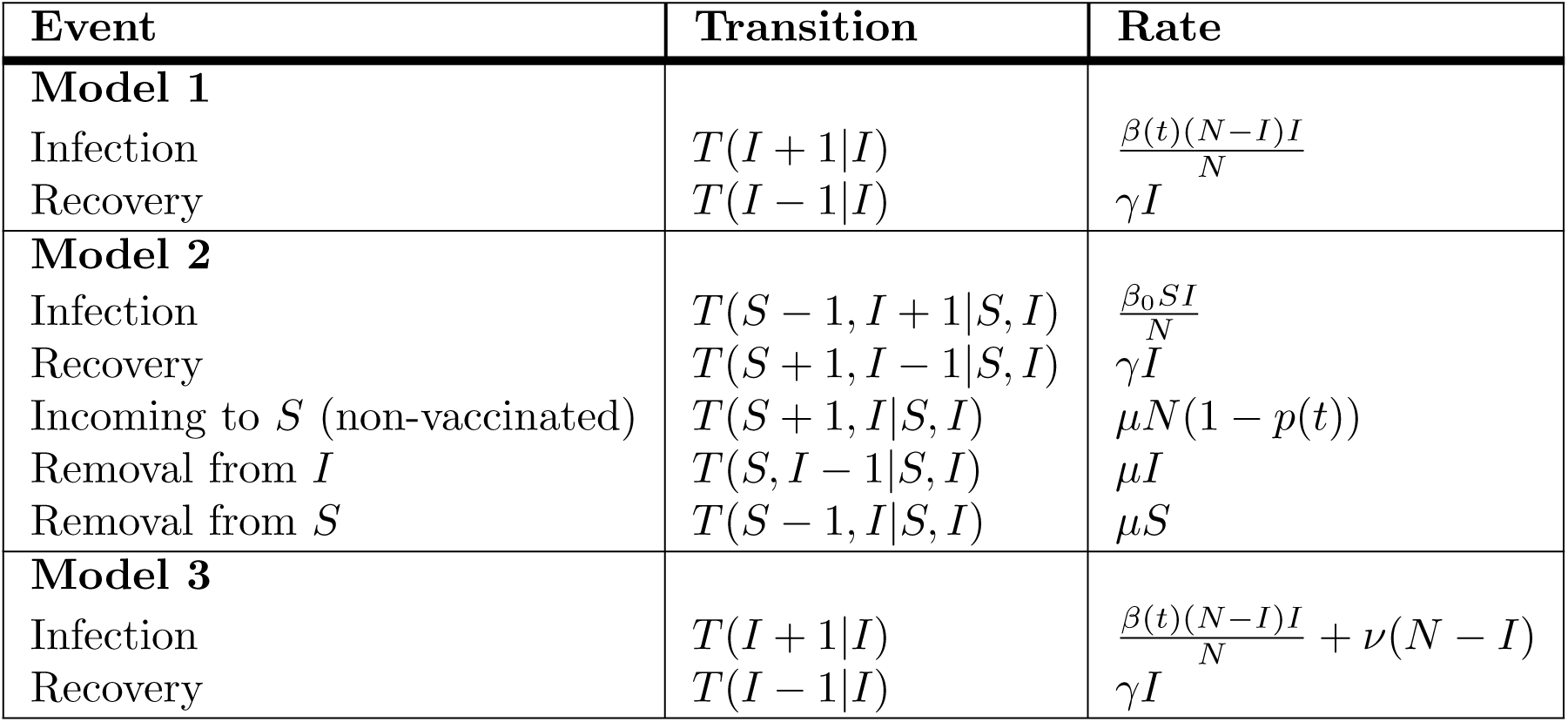
Transition probabilities in prevalence theory for all models.

where *β*(*t*) changes slowly in time, given by,

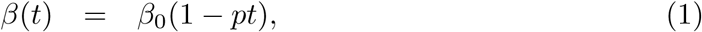

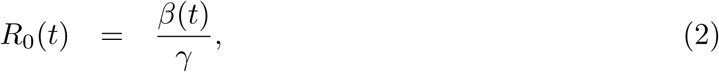

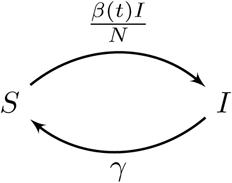

and we fix the population such that *N* = *S* + *I*. Previously work has shown that the fluctuations, *ζ*, about the prevalence steady state, 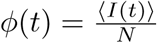 can be separated using the linear noise approximation [28]. The corresponding SDE for *ζ* defined for the SIS is given by [8],

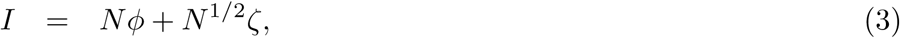

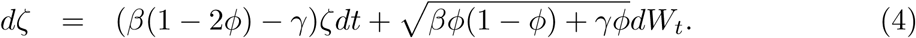

### Model 2: SIS with increasing vaccination coverage (births and deaths)

We consider an SIS model where a proportion of susceptible individuals are vaccinated and gain immunity to the disease. By increasing the proportion of individuals vaccinated *p*(*t*), this control will reduce the effective reproduction number as the susceptible populations is depleting. Births and deaths are considered to allow for a non-zero steady state of *I* initially, and to ensure that the susceptible population does not decrease to zero. By increasing the proportion of individuals vaccinated *p*(*t*), the system is pushed away from this steady state. The remaining (unvaccinated) individuals enter the susceptible component as shown below,

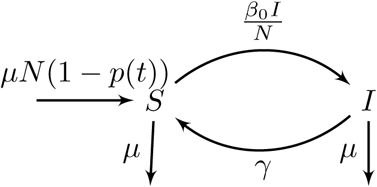

and we gradually increase the proportion of vaccinated individuals over time by,

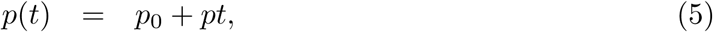

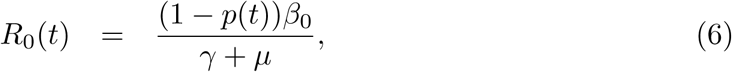

to push the system through the critical transition at *R*_0_ = 1. We interpret the dynamics of the fluctuations of these SDEs with a two-dimensional Fokker-Planck Equation (see supplementary text S1 Appendix and Table 1 for transition rates),

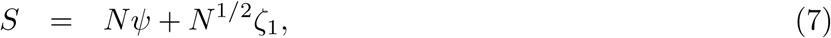

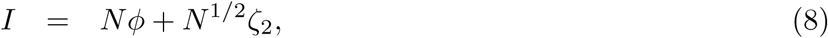

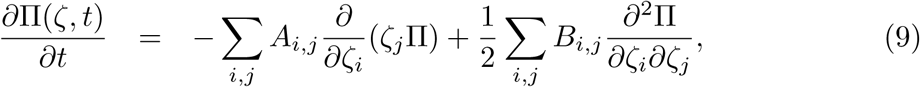

where *ζ*_1_ defines the fluctuations about the susceptibles 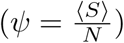 and *ζ*_2_ defines the fluctuations about the infecteds 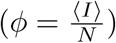, and,

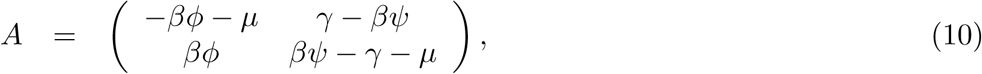

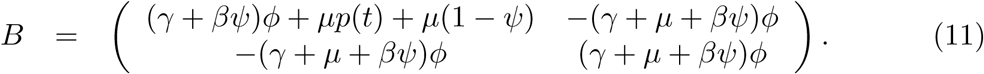

In both cases it is assumed that the fluctuations can be separated linearly from the steady state. Many statistics of this system can be described by the covariance matrix Θ, given by Θ_*ij*_⟨⟨*ζ*_*i*_*ζ*_*j*_⟩⟩ = ⟨*ζ*_*i*_*ζ*_*j*_⟩ − ⟨*ζ*_*i*_⟩⟨*ζ*_*j*_⟩ which satisfies,

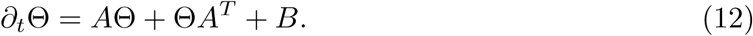

In particular, the variance of the fluctuations about the infectious steady state is given by Θ_22_.

### Model 3: SIS with external force of infection and increasing transmission

Finally we consider the SIS model with external infection which has been used to investigate EWS in prevalence and in incidence [4, 5]. We demonstrate how our analytical results compare for this system, and illustrate differences when applied to disease elimination.

In this model, in addition to the underlying SIS dynamics, susceptible individuals can be infected by an external force of infection (governed by parameter *ν*) that does not depend on the level of infection, shown in the schematic below,

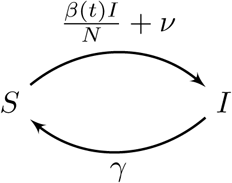

and we consider the model in a stochastic formulation, with transition probabilities given in Table 1. Disease emergence is driven by increasing the effective contact rate *β*(*t*) over time, that slowly increases *R*_0_ through the critical transition at *R*_0_ = 1,

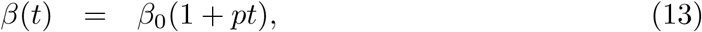

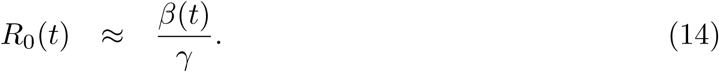

The fluctuations, *ζ* about the steady state are an extension of those in Model 1 to include the external force of infection and satisfy,

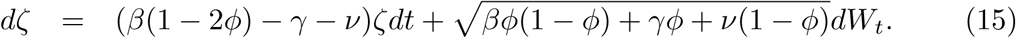

The study by O’Dea *et al.* approximated an SIS with a Birth-Death-Immigration process, where an immigration event approximates the external force of infection; birth events give new infections and the death component is analogous with recovery events. O’Dea *et al.* derive statistics for incidence data by monitoring the number of individuals recovering (e.g. the transition rate *T* (*I −*1*|I*) = *γI*). Results from this study can be found in the supplementary text (S1 Appendix).

One limitation of this methodology is its inability to extend to other systems. This derivation would need to be computed again for each specific example. In particular, if one wanted to consider a simpler system with no external forcing, by setting *ν* = 0, it would make this result redundant - prompting our study for generic EWS that can describe all systems.

## Incidence Theory

A counting process can be used as a generalised theory to understand the dynamics of the number of new events over a period of time. In particular, a diverse range of data types can be described by a counting process and this motivates us to characterise how statistics of such processes behave on the approach to a critical transition. Incidence (the number of new cases, *N*_*t*_) is a counting process, which is known to be described by a non-homogeneous Poisson process {*N* (*t*) : *t ∈* [0, *∞*)} with time dependent rate *λ*(*t*),

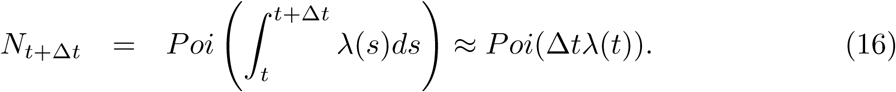

where the integral approximation holds for Δ*t* sufficiently small. In the supporting text (S2 Fig) we demonstrate that for our parameters, this approximation works well for Δ*t* up to 3. We can derive EWS in disease incidence aggregated over a time interval Δ*t* (e.g. daily, weekly, biweekly cases) using the well-known central moments of the Poisson distribution:

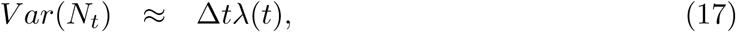

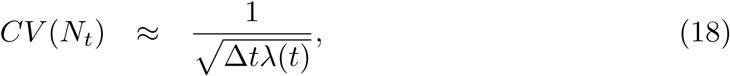

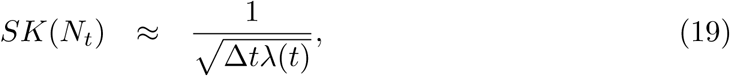

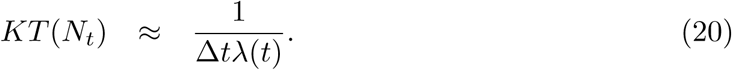

Prior work from O’Dea *et al.* [4] has also incorporated under-reporting using a negative binomial distribution; this can be included in this model when the rate *λ*(*t*) is gamma distributed.

Without under-reporting the rate of new cases is given by the incoming transition probabilities to the infectious state,

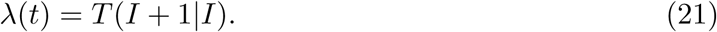

A common form of this force of infection is,

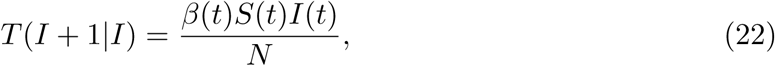

so that *λ*(*t*) depends on the prevalence of infection, *I*(*t*). For Model 1, *β*(*t*) is a function of time whereas in Model 2 *β*(*t*) = *β*_0_ is fixed. Infection can also be increased in other ways such as an external force of infection (Model 3), 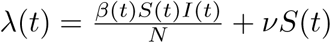, that is typically used to describe zoonotic spillover events or as an approximation for human migration.

## Rate of Incidence Theory

We consider the rate of incidence (or the rate of the Poisson process) *λ*(*t*) = *T* (*I* + 1*|I*), which can be described dynamically with an SDE. Our analyses shows that the critical transition of the rate of the Poisson process corresponds to prevalence models (e.g. at *R*_0_ = 1) and importantly exhibits behaviours predicted by CSD.

We investigate here calculating statistics on the rate of incidence (RoI) and its potential to be used as an EWS for disease transitions.

Below we present our analytical results describing statistical indicators for each model. These theoretical solutions can be used to derive time-varying indicators for the fluctuations of the rate of incidence. Full derivations of the analytical work are given in the supporting text: S1 Appendix.

### Model 1

When considering Model 1 (SIS with decreasing transmission rate), we describe the change in the rate of incidence as 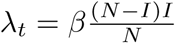. In particular, by considering the time derivative of *λ*_*t*_ we can conclude that the fixed points of the rate of incidence can be described by the transcritical bifurcation at *R*_0_ = 1. We find that the stability of the fixed points of *λ*_*t*_ also correspond to those of *I*, as expected.

We describe the fluctuations, *ω*, about the steady state of 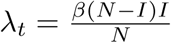 using the linear noise approximation (LNA). We are interested in statistics calculated on the fluctuations about the rate of incidence, to develop new indicators of disease elimination. We derive the resulting analytical solution for *ω* using Ito’s Change of variable formulae (details in supporting text: S1 Appendix) to approximate *ω* with the following Gaussian process:

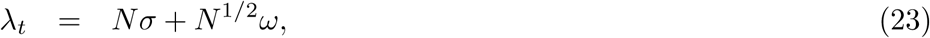

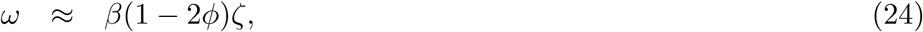

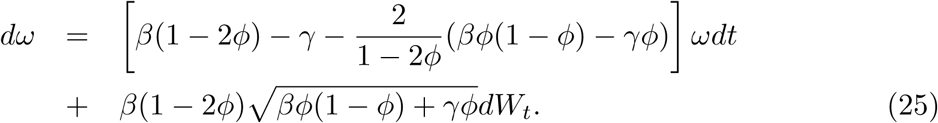

In particular, the changing behaviour of the variance of the rate of incidence as the system approaches disease elimination can be calculated from the SDE Eqn, 25,

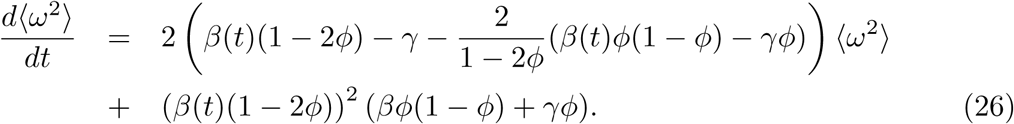

### Model 2

If we consider models where there is population-level immunity, then 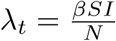 and we can no longer reduce the dimension of incoming transitions using *S* = *N−I*. This can be seen in Model 2 (SIS with increasing rate of vaccination), in particular the prevalence analysis of these systems presented in the Methods Section results in a multivariable Fokker-Plank Equation.

However, we can similarly describe the fluctuations, *ω*, about the steady state of 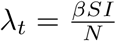 using the linear noise approximation (LNA) as in Model 1.

We again use Ito’s Change of variable formulae for the multivariable system (which will depend on *ζ*_1_ and *ζ*_2_) to approximate *ω*. This leads to an SDE equation which depends on the description of *ζ*_1_ and *ζ*_2_ (eqn. 9). In particular, we are interested in statistics of the rate of incidence, such as the variance, which can be simplified in terms of the original covariance matrix Θ (eqn. 12) and mean-field equations of infectious (*φ*) and susceptible (*ψ*) individuals,

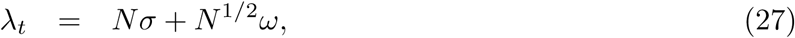

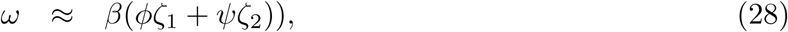

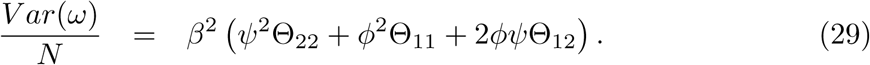

### Model 3

Incorporating an external force of infection into the SIS model is a direct extension of the Model 1 analysis on the rate of incidence. Incoming transitions into the infectious class increases from 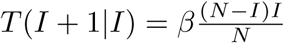 to 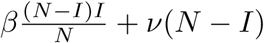, the case which is observed in Model 3 (SIS with external force of infection and increasing transmission).

Following, we can describe the fluctuations about the rate of incidence (*ω*) using the LNA. In particular,

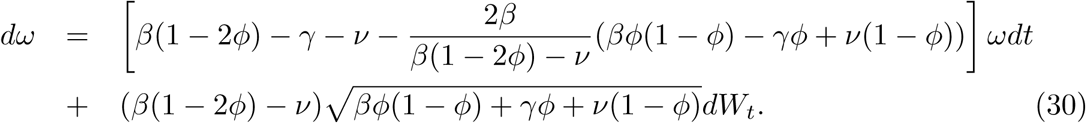

As previously, we can derive statistics of *ω* from the solution of this SDE. In particular, since the SDE is linear in *ω* then we can describe *ω* as a Gaussian variable with mean zero and variance given by the solution to the following ODE,

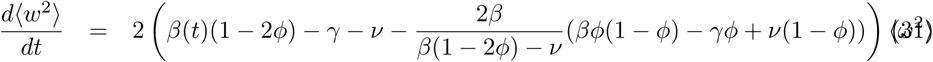

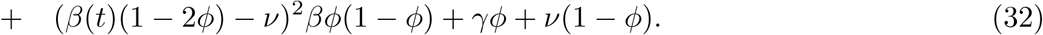

## Simulated Study

We use the Gillespie algorithm to simulate each model, using time varying parameters (*β*(*t*) for Model 1 & 3 and *p*(*t*) for Model 2) to drive the model either to extinction (Model 1 & 2) or emergence (Model 3). We record prevalence outputs at time steps of 0.1 per day and we aggregate incidence outputs to daily time steps Δ*t* = 1. Parameters common to each model are given in table 2. In Model 1, the transmission parameter *β* was reduced from *β*_0_ = 1 to 0, slowly forcing *R*_0_ = 5 to 0. In Model 2, the rate of vaccination was increased from *p*_0_ = 0 to 1, slowly forcing *R*_0_ = 5 to 0. In Model 3, the transmission parameter *β* was increased from *β*_0_ = 0.12 to 0.24 so that the basic reproduction number increases from *R*_0_*≈*0.6 to*≈*1.2.

**Table 2.** Parameter values shared among all models.

| Parameter | Description | Value |
| --- | --- | --- |
| $\beta_0$ | Initial Transmission Rate | $\beta_0^{\{1,2\}} = 1, \beta_0^{\{3\}} = 0.12$ |
| $\gamma$ | Recovery rate | $\gamma^{\{1,3\}} = 0.2, \gamma^{\{2\}} = 0.18$ |
| $\mu$ | Population turn over rate | $\mu^{\{2\}} = 0.02$ |
| $p_0$ | Initial vaccination rate | $p_0^{\{2\}} = 0$ |
| $p$ | Rate of change of $\beta_0$ or $p_0$ | $p^{\{1,2,3\}} = 0.002$ |
| $\nu$ | External rate of infection | $\nu^{\{3\}} = 0.001$ |
| $N$ | Population Size | $N = 2000$ |
| $\Delta t$ | Time aggregation of incidence data | $\Delta t = 1$, daily |
| $T$ | Time simulations run for | $T = 500$ (after burn in of 300 days) |
| Ext | Simulations with reducing parameters inducing extinction (Model 1 & 2) |  |
| Emg | Simulations with increasing parameters inducing emergence (Model 3) |  |
| Fix | Simulations with fixed parameters (null) |  |

Table notes: values in braces directs to the model number which was implemented at that value. Parameters without braces are shared amongst all models.

A drawback of using the rate of incidence (RoI) as a measure of disease elimination, is the need to develop methods to extract this rate from incidence data. In our simulation study, we calculate the RoI in two ways. Firstly, using simulations of prevalence and taking the product 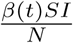. This method, although unrealistic as it requires knowlege of prevalence (*I*), demonstrates the accuracy of the analytical results, as it is the “true” definition of RoI. An alternative method uses the Poisson property of incidence, illustrating that the rate of incidence *λ*(*t*) also corresponds to the mean incidence over time. Therefore, our second method evaluates RoI by linearly smoothing the Gillespie output of incidence (*N*_*t*_) on a rolling window, giving the approximated mean number of new cases for each realisation (we refer to this method as “approximated” RoI). We can then calculate statistics of the fluctuations of the rate of incidence over multiple realisations of this smoothed time series.

For each model, we also perform simulations where the disease has not fully gone through a critical transition (null model) which we refer to as Fix simulations. This null model has no control mechanism and the disease fluctuates about the fixed endemic steady state, at *R*_0_ = 5 (Model 1 & 2) and *R*_0_ = 0.5 (Model 3).

Before calculating the time changing statistics, we detrend each simulation by removing the mean over all realisations of that setting (Ext, Emg or Fix). We are interested in five common statistical indicators: variance (V), coefficient of variation (CV), skewness (SK), kurtosis (K) and autocorrelation lag-1 (AC(1)). We illustrate how EWS change over time, and how accurate the theory is to predicting these trends. In particular, we are interested in the trends over time and whether these time series properties of EWS are the same for prevalence and incidence data.

The Kendall-tau score gives a measure of an increasing or decreasing trend of each statistic over the time series (where *N* is the number of time points). We use the measure to evaluate whether a statistic corresponds to an increasing or decreasing trend and compare this for different data types (prevalence, incidence and RoI). The Kendall-tau score is defined as [25],

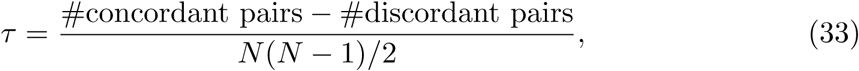

where two points in the time series 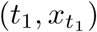 and 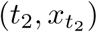 with *t*_1_ *< t*_2_ are said to be a concordant pair if 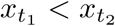, and a discordant pair if 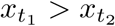. If the two points are equal 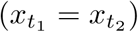 then the pair is neither concordant or discordant.

We calculate each statistic on a moving window (size 50) for each detrended simulation, and then evaluate the Kendall-tau score. We compare the Kendall-tau scores calculated on simulations going through a critical transition with null simulations, and we then calculate receiver operating characteristic (ROC) curves by considering the null model to be negative and the other models to be positive. We compare the performance of each model statistic using the area under the curve (AUC). Good statistics have an AUC close to 1 or 0 since this indicates the statistic is far from picking by chance.

## Results

### Variance

Variance is one of the most intuitive statistical indicators. As a system approaches a critical transition the time taken to recover from small perturbations increases, as described by Critical Slowing Down theory. This can be observed in the fluctuations about the steady state, which on the approach to a critical transition take longer to return and consequently vary far more, defining the increasing nature of variance as an early warning signal.

We evaluate analytical solutions of the variance in prevalence using the derived SDE for each model (Model 1: Eqn.4, Model 2: Eqn.9, Model 3: Eqn.15). We compare this to theoretical solutions of the variance in incidence, using the transition rates for each model (Table 1) to compute the rate of the Poisson process *λ*. Figure 1 presents the simulated statistics for both prevalence and incidence theories of elimination and emergence, where we have plotted the variance between multiple homogeneous simulations at each time point (described in supporting text: S3 Fig).

**Fig 1.**
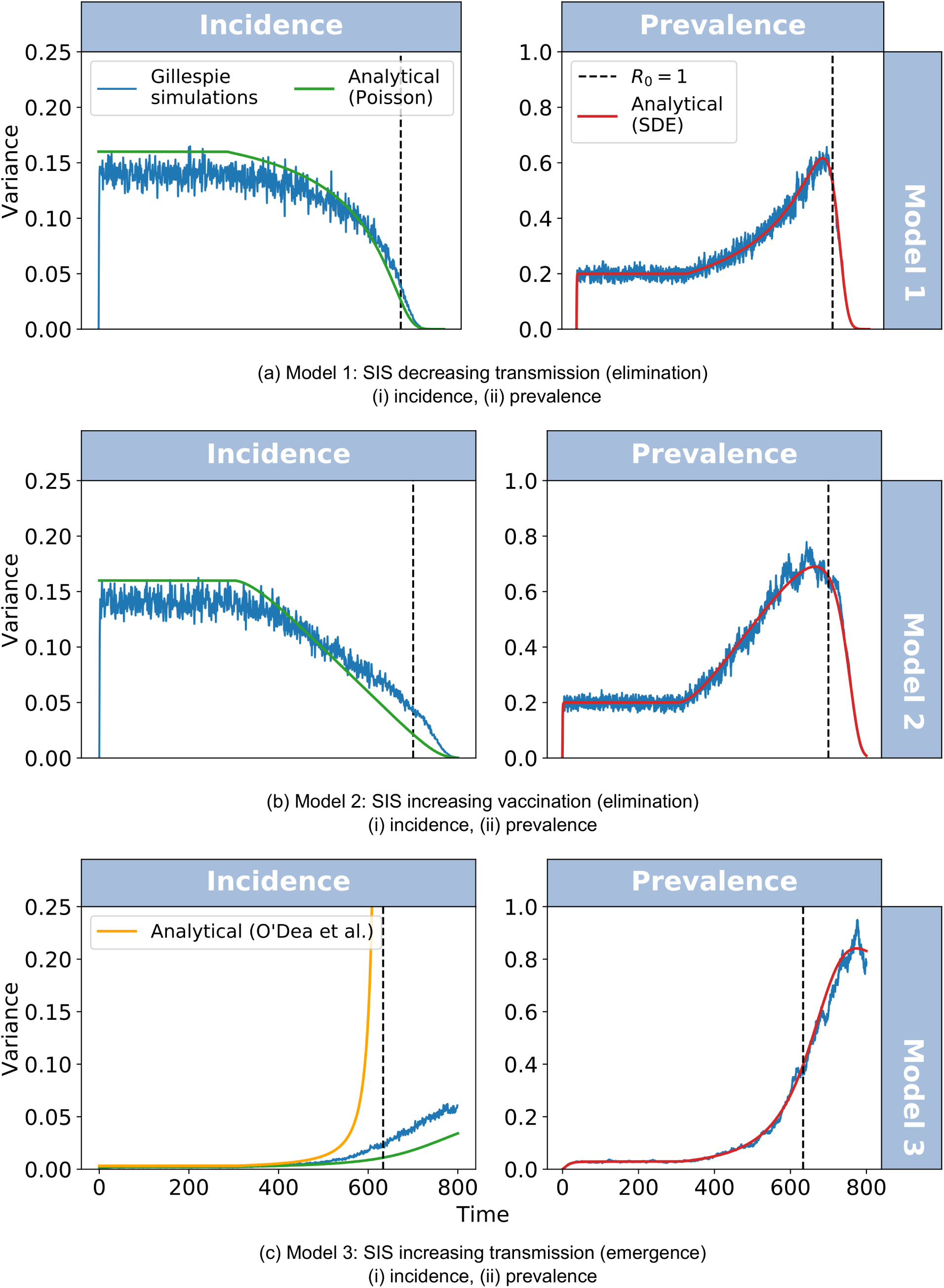
Comparing predictions to simulations for variance. For each model (Model 1: SIS decreasing transmission; Model 2: SIS increasing vaccination; Model 3: SIS increasing transmission, emergence) we calculate the variance between 500 homogeneous realisations at every time step (daily). Each figure shows: Poisson process distribution (green line); dynamic predictions (red line) and Gillespie simulations (Ext: Model 1 and 2 and Emg: Model 3, blue line). The last model also shows the dynamical prediction from O’Dea *et al.* which was derived for this specific system (orange line).

Our prediction for the variance is similar to the stochastic simulations with a slight underdispersion in the incidence simulations (Figure 1 a(i), b(i) and c(i)), since the theory (rate of Poisson process, green line) is higher than that of the variance in the fluctuations of the simulations (blue line). We note that our solution for Model 3 (Fig. 1) follows the gradient of the stochastic simulations more closely. Since the analytical solution by O’Dea *et al.* (orange line) is evaluated at the steady state then the result diverges at the critical transition. It can be shown that for larger values of *β*_0_, O’Dea *et al.* results fit closer to the stochastic simulation, although the general trend of variance for both approaches follows the simulations.

We observe that variance in prevalence simulations (Figure 1 a(ii), b(ii) and c(ii)) increases on the approach to the critical transition, as predicted by critical slowing down. In comparison the variance in incidence decreases before the critical transition for all disease elimination models (Model 1 Fig. 1 a(i) and Model 2 Fig. 1 b(i)) and increases similarly to prevalence for the disease emergence model (Model 4 Fig. 1 c(i)). This is contradictory to the theory of critical slowing down theory which predicts that derivations from the steady state values return increasingly slowly on the approach to a transition.

As expected by our Poisson process analysis, the variance of this system should be the same as the mean of the system. Therefore for disease elimination models, we should expect a decreasing variance (along with a decreasing mean) when calculated on incidence data, in contrast to an increasing variance with prevalence data. Likewise with disease emergence models we expect an increasing variance to correspond to the increasing mean. This demonstrates that our analysis of incidence has successfully predicted the time-varying variance for these different systems.

## Rate of incidence

We have observed that incidence data does not approach a critical transition as described by critical slowing down theory. Consequently we demonstrated in Figure 1 that the variance of incidence does not necessarily increase on the approach to a critical transition. A new approach for working with incidence-type data is to consider the rate of incidence, *λ*(*t*) = *T* (*I* + 1*|I*), which for each model we have derived the dynamical SDE (see Methods).

### Variance (RoI)

The analytical variance in the rate of incidence is presented in Fig. 2 (orange line). We find that the theoretical analysis supports the simulation studies and provides us with an understanding of how statistical indicators calculated in RoI data change on the approach to a critical transition. We observe an increasing variance in the rate of incidence before a critical transition; a time series trend which if exhibited in real-world data could be used to anticipate disease tipping points.

**Fig 2.**
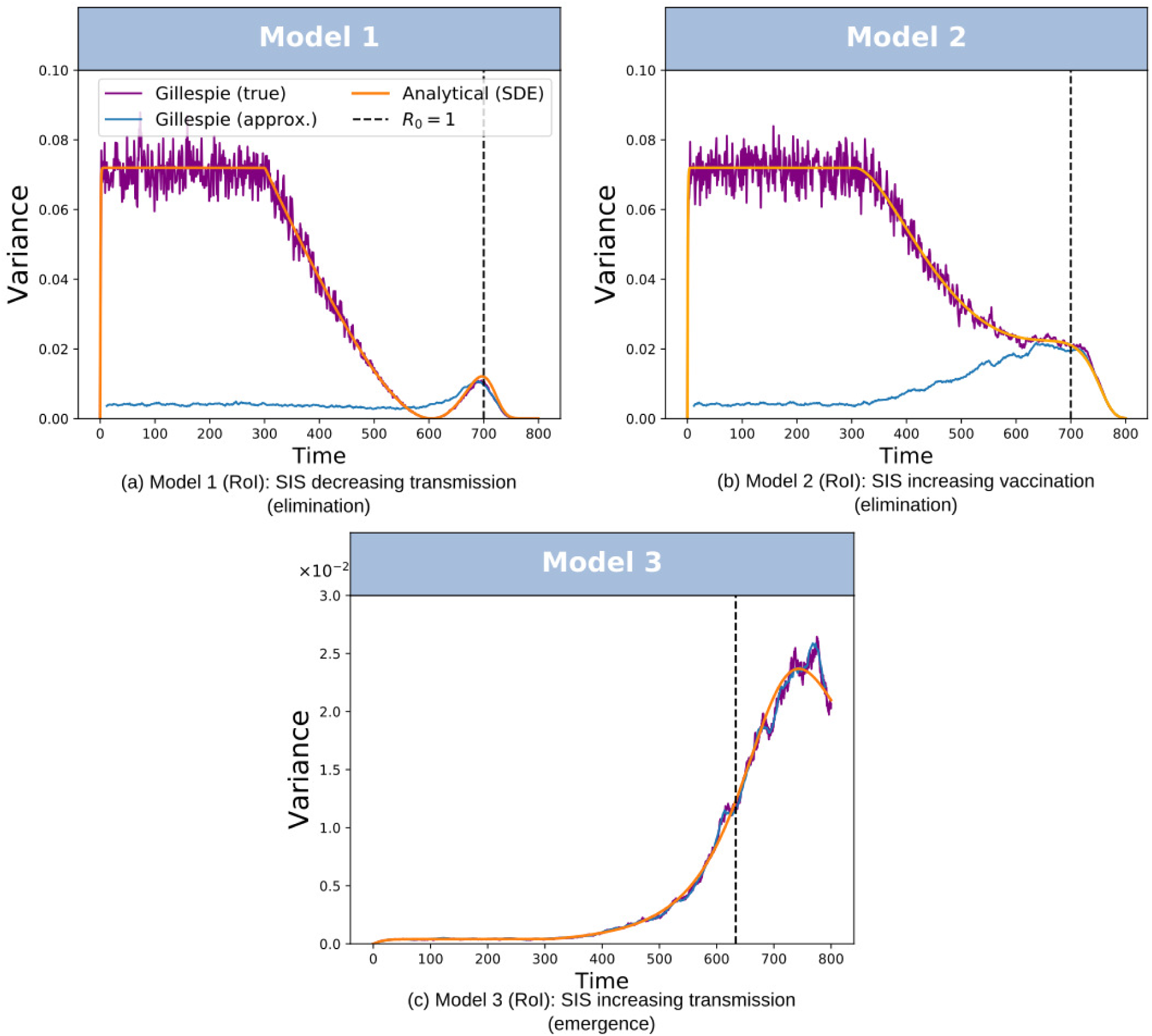
Variance calculated on the rate of incidence. For each model (Model 1: SIS decreasing transmission; Model 2: SIS increasing vaccination; Model 3: SIS increasing transmission, emergence) we calculate the variance on the rate of incidence. between 50 homogeneous realisations at every time step (daily). Each figure shows: dynamic solution (orange line); rate calculated from Gillespie simulations of incidence after smoothing (light blue line) and Gillespie simulations prevalence multiplied by time-varying susceptible and effective contact rate (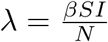, purple line).

We present results calculated in RoI simulations using the two methods: “true” and “approximated” RoI, in Fig. 2. The first method uses prevalence data (“true”, purple line) and corresponds well with the analytical solution (orange line) for all models and the latter method (smoothing incidence data “approximated”, blue line) fits particularly well for Model 3 (Fig. 2(c)). However it does not follow as closely to some time-varying properties of the variance for Model 1 & 2 (Fig. 2(a) and (b)) respectively. Although the early dynamics are misrepresented for Models 1 and 2, all time series indicate an increasing variance on the approach to the critical transition. In particular, “approximated” RoI can be implemented in practice from incidence type data (blue line), captures this property in all models.

In Fig. 2(a) and (b) we observe that the analytical prediction fits well with the stochastic simulations of 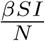 (purple line) for Model 1 and Model 2 respectively. This demonstrates that this theory approximates the behaviour of the system well. Indeed, we observe that approximating the rate of incidence by smoothing Gillespie simulations of new cases and then calculating the variance of this quantity (“approximated” RoI) predicts a similar increasing behaviour before the critical transition, shown in Fig. 2(a) (blue line). This corresponds to the same peak as the analytical prediction and “true” simulations. However, it fails to capture the magnitude of the behaviour earlier on in the dynamics.

An area that still needs to be addressed with this methodology of smoothing new case data is determining a suitable window size. This could result to misleading EWS when used in practice. In the supporting text S1 Fig, we demonstrate that if the disease is approaching elimination at a slower rate, both methods (“true” and “approximated”) converge to the analytical solution. We chose parameters such that Model 1 approaches disease elimination at the same rate as Model 3 approaches disease emergence (*R*_0_ changes from 1.2 to 0, 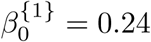). As the system changes slowly enough then the system will be approximately ergodic, such that the moving average resembles the mean incidence. Thus the “approximated” method will be closer to the “true” solution. In comparison, the faster a system changes over time, will correspond to a wider range in incidence cases across the moving window. Resulting in a lower mean over the window which can be seen in Fig. 2(a),(b); although the statistic will be more pronounced at the threshold.

We find that for Model 2 (Fig. 2(b)) the general trend of the variance is less pronounced at the critical transition than observed for Model 1. We observe that the analytical solution (Fig. 2(b) orange line) and true stochastic simulations (Fig. 2(b) purple) only slightly increase before the critical transition, implying this trend would be difficult to detect in real-world data. In particular, the Kendall-tau score which can be an indication of an increasing trend, is negative (decreasing, *τ* =*−*1) for Model 2, whilst for Model 1 and 3 we find that *τ* = 0.987 and *τ* = 1 respectively. Although, we observe that the “approximated” simulations of the rate of incidence (Fig. 2(b) blue line) exhibit similar properties as Model 1. We observe that the early stage dynamics of this method have not predicted the expected behaviour of the analytical solution. It can be noted that *R*_0_ decreases at the same rate as Model 1, suggesting that this can be a result of the approximation when *R*_0_ is not slowly changing. Due to this approximation, it can be observed that the variance of new cases does therefore increase before the critical transition (blue line).

In Fig. 2(c) we observe that both measurements of the variance of *λ*_*t*_ calculated on stochastic simulations of Model 3 have closely followed the analytical solution of variance. As expected the true stochastic simulations (Fig. 2(c) purple line) follow closely to the theory, supporting that this derivation of *ω* is correct. More interestingly, calculating the variance of the rate of incidence directly from simulations of new cases (*N*_*t*_, Fig. 2(c) blue line) has performed far better than when presented in Model 1 (Fig. 2(a)). For Model 3, we observe that the variance of the rate of incidence increases before the critical threshold, where the infectious disease emerges with outbreaking dynamics similar to prevalence for this model. We further found that the early dynamics of the “approximated” RoI simulations represent the true behaviour of the variance. This result may be due to *R*_0_ increasing more slowly in Model 3 than the rate it decreases at in Model 1, satisfying the ergodic condition.

## Other statistical indicators

In this section, we investigate the potential of identifying an epidemiological transition using five commonly implemented early-warning signals: variance, coefficient of variation (CV), skewness, kurtosis and lag-1 autocorrelation (AC(1)). Exploration of each EWS follows similarly to variance, as analysed above theoretically and numerically for prevalence, incidence and rate of incidence. In the supporting text, time series trends for each indicator are presented for each dataset and model (S3 Fig: variance, S4 Fig: CV, S5 Fig: Skewness, S6 Fig: kurtosis, S7 Fig: AC(1)), along with analytical analyses for these indicators (S1 Appendix).

Here, we quantify these time series trends for each statistical indicator using the Kendall-Tau score as a measure of an overall increasing or decreasing trend. We present in Fig. 3 the predictive power of each statistical indicator using its time-changing trend to classify simulations as either extinct (Ext simulations), emerging (Emg simulations) or null simulations (Fix simulations). We calculate the area under the ROC curve (AUC) by comparing the Kendall-tau score calculated over each time series up to two end points: before the critical transition (*t*_1_) and after the critical transition (*t*_2_) which gives an overall score of the true/false positive rate.

**Fig 3.**
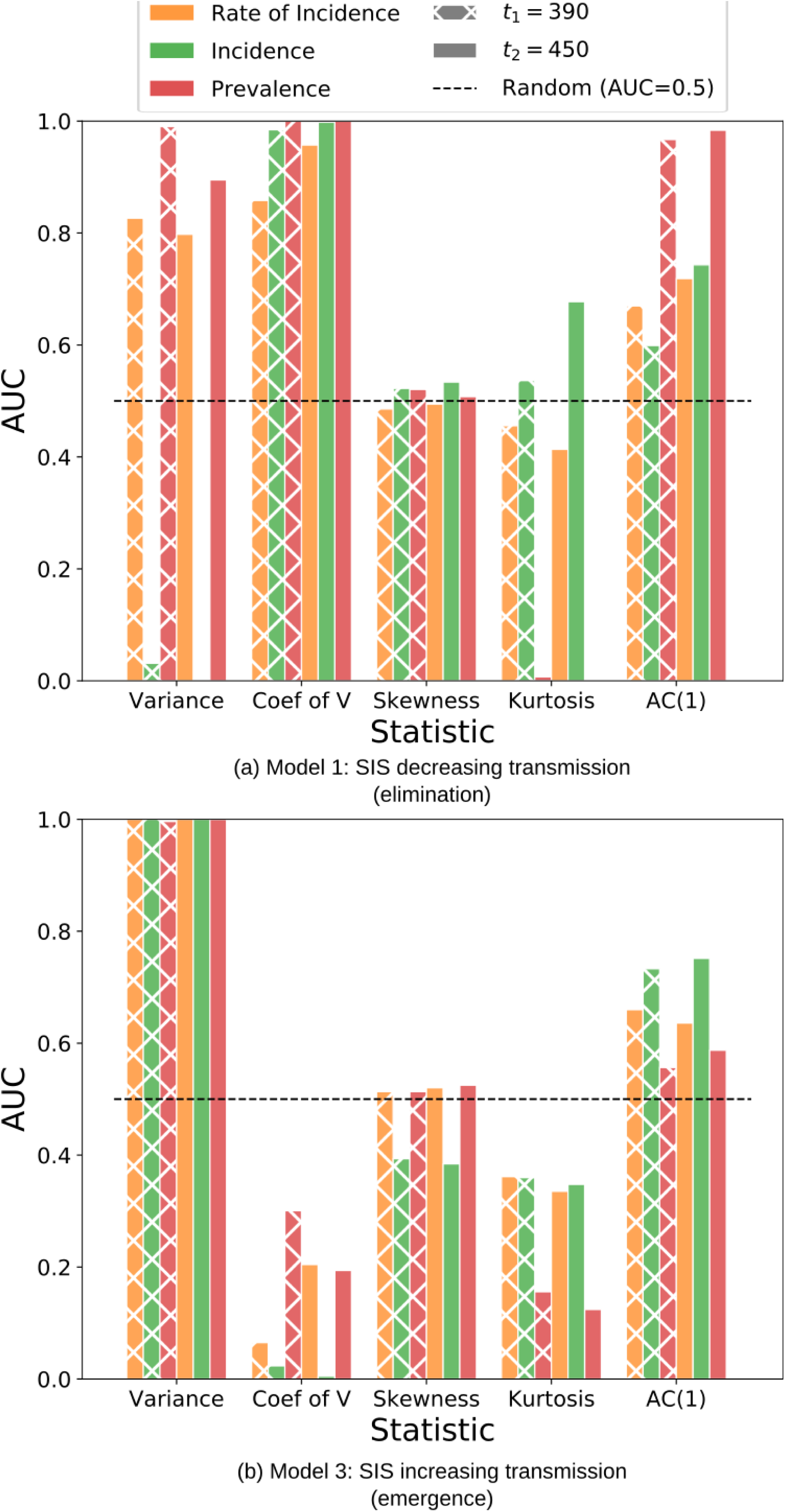
AUC scores for different EWS. We compare the performance of 5 common statistical indicators for Model 1 (a) for disease elimination and Model 3 (b) for disease emergence. For each ROC curve, we measured the AUC which is an indication of how predictive each indicator is by its ability to distinguish between elimination simulations and the null model. A score closer to 0.5 signifies the worst performance (random diagnosis). We evaluate the Kendall-tau score up to before the critical transition (*t*_1_ = 390) and after the critical transition (*t*_2_ = 450), which gives an indication if the EWS is increasing or decreasing. A score of 1 demonstrates that it is possible to identify all Ext simulations when compared to null simulations by its increasing trend (i.e. perfect sensitivity, true positive rate). A score of 0 means that there is zero sensitivity and instead the simulations are decreasing.

The AUC score gives a predictive measure between different indicators, which we use to assess their performances. The closer the AUC is to 0.5 signifies the worst the statistic’s performance at anticipating a critical transition. This is analogous with randomly selecting simulations that are the null and disease elimination (Model 1, Fig. 3(a)) or disease emergence (Model 3, Fig. 3(b)). In particular, skewness is a poor indicator because of its inability to identify disease elimination with any type of disease data it is applied to (rate of incidence, incidence and prevalence). Identifying emergence with skewness in prevalence or RoI data (red and orange bars respectively) is also very poor and its predictive ability is only slightly increased with incidence (green bars).

A score close to 1 indicates nearly perfect sensitivity and specificity. For each EWS, we assume that an increasing trend represents a disease going through a critical transition. As a result a AUC score of 1 informs us that the indicator is increasing and that it is possible to identify all Ext/Emg simulations when compared to the null simulations by its increasing trend. Notably, in Fig. 3 coefficient of variation calculated on all types of disease data (rate of incidence, incidence and prevalence) and for both Model 1 & 2, exhibits a near perfect ability to identify the increasing trend.

An AUC score of 0 demonstrates that the time series trend is instead decreasing and as such it doesn’t correspond to the predetermined prediction. A perfectly diagnosed decreasing indicator when compared to the null model will result in zero sensitivity under these conditions and an AUC score of 0. Fig. 3 highlights which indicators are in some cases increasing (AUC close to one), decreasing (AUC close to zero) or are poor indicators (AUC close to 0.5). In particular, as discussed in the previous section, variance always increases prior to disease emergence (Fig. 3(b)). However, for disease elimination (Model 1: Fig. 3(a) and Model 2: S11 Fig) results are substantially different when we compare variance calculated in rate of incidence and prevalence (orange and red bars respectively) with incidence (green bars). For RoI and prevalence data types, the statistical signature is an increasing variance with an AUC near 1. This is in contrast to the latter where the trend is decreasing with an AUC near 0. However, the results for variance (both increasing and decreasing) are highly predictive (*|AUC −* 0.5*| ≈* 0.5). Thus, if a system is not known or there is difficultly in determining the type of data, incorrect conclusions could be drawn when interpreting the time series trend.

## Discussion

While studies for EWS on incidence-type data have been growing in recent years, theoretical exploration of how these indicators change on the approach to a critical transition have been neglected. In this paper, we have shown that the typical trends of EWS that precede a critical transition are exhibited in prevalence-type data but do not always exist in incidence-type data. In particular, we have focused our investigation on the trend of variance over time as an infectious disease system approaches a tipping point.

Prior work has shown that variance in incidence increases on the approach towards disease emergence. However, our work highlights that this property is not a result from critical slowing down theory as first expected. We have shown it is a consequence of the counting process that can approximate incidence-type data. As such, we demonstrated that the variance in incidence is expected to follow the mean in incidence. In particular, the variance will increase on the approach to disease emergence, but will notably decrease before a disease elimination threshold. We applied these findings to two systems of disease elimination and verified that variance of incidence exhibits a decreasing trend on the approach, following the behaviour of the mean and contradicting critical slowing down theory.

Therefore, it is highly recommended to understand analytically how EWS change on the approach to a critical transition in order to avoid misleading results. The generalised theory of a counting process can be applied to many other systems outside of the scope of epidemiology where we would expect a decreasing variance preceding a critical transition. Potential applications include the observation of animals through camera traps, disease surveillance sampling in wildlife or movements in stock prices, which are all examples of incidence-type data. Notably, a substantial number of studies on ecosystem data, climate data and financial data have observed inconsistencies in statistical indicators [16, 22, 23, 29]. Although we found the Poisson process to be overdispersed in the context of epidemiology, it provides a broad framework which can be extended to many other infectious disease systems using the incoming transition probabilities into the infectious class.

We proposed extracting the rate of incidence (RoI) or intensity of Poisson process from incidence-type data to illustrate that to utilising CSD, such as observing an increasing variance, requires suitable data which undergoes a bifurcation. In particular, we have shown that the critical threshold in the RoI corresponds with that of prevalence; and as expected we demonstrated that the trend in variance in RoI does increase before an imminent epidemiological transition.

We applied five early warning signals to simulated datasets comprising of the three discussed types: prevalence, incidence and rate of incidence. The simulated data we have investigated represents perfect reporting or the “best case scenario”. Often is the case that there is underreporting that may reduce the detectability of signals in real-world data. The work we have presented here can be extended to include a gamma distributed intensity *λ*. Using a gamma distributed rate of incidence will account for reporting errors as described by O’Dea *et al.*

Overall, our study suggests that a robust indicator is one that shares a highly predictive time series trait (*|AUC −* 0.5*| ≈* 0.5). amongst all three data types, even with inconsistent trends (increasing or decreasing). Therefore, we suggest that variance and coefficient of variation are overall good indicators due to their high predictive power in all cases. Coefficient of variation is a robust indicator for disease elimination since the trend is similar between different types of data (Fig. 3 (a)) and S11 Fig. However discrepancies are demonstrated when considering opposite disease thresholds as shown with disease emergence (Fig. 3(b)) which has a decreasing trend for CV and performs less well with disease prevalence data.

However, we found that kurtosis and AC(1) are not robust indicators. Although kurtosis and AC(1) have a predictive trend with prevalence data, this is not typically the data which is readily available. In particular, kurtosis is highly predictive (with a decreasing trend) in prevalence data on the approach to disease elimination (Fig. 3(a)) and fairly predictive with an decreasing trend in the case of prevalence with emergence (Fig. 3(b)); it is a poor indicator for all other types of data. Likewise, although AC(1) has a clear increasing trend for prevalence data elimination systems (Fig. 3(a), S11 Fig), it is less predictive trend for incidence and RoI data. Additionally, the trend is not distinct for any datasets when considering an emergence transition, therefore there is a potential for this indicator to be used incorrectly. In the cases where an EWS is poor in some types of data but good for others could lead to misleading judgements of systems, and therefore are not robust.

These findings support prior work on prevalence and initial work from O’Dea *et al.* and Brett *et al.* with incidence-type data. Our analytical exploration of incidence has indicated a new data source, RoI, which can be extracted from incidence timeseries. A potential powerful tool would be to compute variance and CV indicators with different types of data (incidence, rate of incidence and prevalence) and ensemble these. An ensemble or combination of multiple statistical indicators was suggested by Drake & Griffen [13] and has been applied to case studies with the same data-type and a combination of EWS by Kefi *et al.* [30] to help interpret between different critical transitions and also has successfully detected transitions using an ensemble of different time series data [12]. This suggests a potential approach to achieve a single metric from a combination of indicators calculated on multiple timeseries data with different trends, such as we have observed with incidence and RoI, to achieve a more pronounced indication of disease transitions.

Additionally, further work would be to include a heterogeneous ensemble as suggested by O’Dea *et al.* [4], whereby all parameters are sampled randomly for each realisation rather than being equal. This will lead to more realistic results, as each parameter sample represents time series data from different locations, as suggested by studies on spatial statistics, a promising method for addressing limited data [8, 17, 18]. Comparatively, we have shown here that computing the statistics on a homogeneous ensemble although unrealistic, it returns exact stochastic behaviours of the system and we used this to verify the simulated study with the theory.

In conclusion, there is a tremendous potential for using early warning signals to provide evidence on our progress towards elimination and inform public health policies. We have indicated that by monitoring simple statistics over time it is possible to observe disease emergence and elimination, which with further development offers a promising solution for an automated system that can update time series statistics in real-time as new data becomes available. This would be particularly useful for emerging diseases where EWS could be used to prompt early detection and help aid rapid responses. The focus of our paper has provided insight on how statistics behave for different types of infectious disease data, where we considered suitable data which could be incorporated into such monitoring system. We have researched the resemblance of observed time series results between different data types, a necessary exploration for the development of EWS before they can impact decision making. We reported that some indicators traits are inconsistent across all data types and some EWS differ significantly between disease thresholds: elimination and emergence. Knowledge of the type of data which has been collected is imperative to avoid misleading judgements in response to time series trends. Our work has provided analytical evidence to understand why results differ, improving our ability to monitor EWS for infectious disease transitions.

## Supporting information

**S1 Appendix Analytical Derivations**

**S1 Fig.Slower rate reduces discrepancies between theory and simulated variance in “aproximated” RoI.**

**S2 Fig.Sensitivity to time aggregation (**Δ*t***) when approximating non-homogeneous Poisson process**

**S3 Fig.Comparing predictions to simulations for variance.**

**S4 Fig.Comparing predictions to simulations for coefficient of variation. S5 Fig.Comparing predictions to simulations for skewness.**

**S6 Fig.Comparing predictions to simulations for kurtosis.**

**S7 Fig.Comparing predictions to simulations for autocorrelation lag 1.**

**S8 Fig.Evaluating the MSE between theoretical prediction and simulations for different window sizes.**

**S9 Fig.ROC curves for indicators V, CV and AC(1).**

**S10 Fig.ROC curves for indicators Skewness and Kurtosis.**

**S11 Fig.AUC scores for different EWS (Model 2).**

